# Induction of Antioxidant and Detoxifying Enzymes via the Nrf2-ARE Pathway by Oriental Bezoar

**DOI:** 10.1101/2021.01.25.428174

**Authors:** Tomoe Tsubonoya, Eiji Inoue, Keiichi Sudo, Yasuharu Shimizu

## Abstract

**Objectives:** Oriental bezoar, a gallstone formed in the gall sac of *Bos taurus* Linné var. *domesticus* Gmelin (*Bovidae*), has been used as a crude drug in oriental medicine for stroke, convulsion, epilepsy, swelling and pain in the throat, and high fever. Despite reports on the antioxidative effect of oriental bezoar-containing drugs, *in vitro*-cultured calculus bovis (a substitute for oriental bezoar), and its components (namely, bilirubin and biliverdin), the intracellular mechanism of the antioxidative action of oriental bezoar remains unclear. Therefore, we studied the effect of oriental bezoar on the nuclear factor erythroid 2-related factor 2 (Nrf2)-antioxidant response element (ARE) pathway, which transcriptionally regulates genes encoding antioxidant and detoxifying enzymes, such as heme oxygenase-1 (*HO-1*), glutathione-S-transferase alpha 1 (*GSTA1*), and quinone oxidoreductase 1 (*NQO1*), to elucidate its antioxidative effect.

**Methods:** ARE promoter activity and mRNA expression (*HO-1, GSTA1*, and *NQO1*) in HepG2 cells supplemented with test drugs were examined *in vitro* by luciferase reporter assay and real-time PCR, respectively.

**Results:** Oriental bezoar induced ARE promoter activity and increased the mRNA expression level of *HO-1, GSTA1*, and *NQO1*. Its components, namely, bilirubin and biliverdin, also increased the ARE promoter activity.

**Conclusion:** These results suggest that oriental bezoar can induce antioxidative and detoxification effects via the Nrf2-ARE pathway.

## Introduction

Oriental bezoar is a gallstone formed in the gall sac of *Bos taurus* Linné var. *domesticus* Gmelin (*Bovidae*), which contains bilirubin, biliverdin, bile acids, and amino acids [1–5]. It has been used as a crude drug in oriental medicine for stroke, convulsion, epilepsy, swelling and pain in the throat, and high fever [1–3, 5]. Some of these applications of oriental bezoar are derive from its antioxidative effects [3]. Based on our previous papers [6, 7], Reiousan®, a crude drug consisting of oriental bezoar and ginseng, has shown antioxidative effects *in vitro* and a protective effect against ischemia–reperfusion injury *in vivo*. In addition, *in vitro*-cultured calculus bovis, a substitute for oriental bezoar, is reported to have antioxidative effects [8, 9]. Bilirubin and biliverdin, which are the components of oriental bezoar, are also reported to have antioxidative effects [10–12]. These reports suggest that the aforementioned use of oriental bezoar can be partially explained by its antioxidative effects. However, the intracellular mechanism of action remains unclear.

The nuclear factor erythroid 2-related factor 2 (Nrf2)-antioxidant response element (ARE) pathway plays a role in the biological defense against oxidative stress and foreign substances [13]. Upon exposure to oxidative stress and foreign substances, Nrf2 translocates from the cytoplasm to the nucleus and binds to the ARE to transcriptionally activate genes encoding antioxidant and detoxifying enzymes, such as heme oxygenase-1 (HO-1), γ-glutamyl cysteine ligase, thioredoxin reductase-1, superoxide dismutase, glutathione peroxidase, catalase, glutathione-S-transferase alpha 1 (GSTA1), quinone oxidoreductase-1 (NQO1), and UDP-glucuronosyltransferase [13]. Although the Nrf2-ARE pathway is vital for maintaining homeostasis [13, 14], Nrf2 levels decline in aging animals [15, 16]. Therefore, activating the Nrf2-ARE pathway has been considered a potential strategy to develop drugs for preventing various diseases caused by oxidative stress and foreign substances [13, 14].

In this study, we studied the effect of oriental bezoar on the Nrf2-ARE pathway *in vitro* to elucidate the intracellular mechanism of its antioxidative effect.

## Materials and Methods

### Cells

HepG2 cells (RIKEN BioResource Research Center, Ibaraki, Japan) were stored in liquid nitrogen. The number of cell passages for gene expression analysis and cytotoxicity assay was 1 to 16, whereas that for reporter assay was 18 to 22.

### Test drugs

Oriental bezoar (Kanai Shoten, Tokyo, Japan and Miyachu, Osaka, Japan) was obtained in Brazil, and it was mixed and pulverized in our manufacturing department (Kyushin Pharmaceuticals, Tokyo, Japan). The powdered oriental bezoar was suspended in 10% dimethyl sulfoxide (FUJIFILM Wako Pure Chemicals, Osaka, Japan) and extracted using an ultrasonic generator (Model 2510, Branson ultrasonics, CT, USA) for 30 min at 25°C. The extract was sterilized by filtration using a 0.2 µm membrane filter (ADVANTEC, Tokyo, Japan).

Bilirubin, R, S-sulforaphane (FUJIFILM Wako Pure Chemicals), biliverdin hydrochloride (Sigma-Aldrich, MO, USA), and hemin (Alfa Aesar, UK) were dissolved in dimethyl sulfoxide.

### Chemicals

Phosphate-buffered saline, RPMI 1640 medium (FUJIFILM Wako Pure Chemicals), fetal bovine serum, 2.5% trypsin (Thermo Fisher Scientific, MA, USA), penicillin-streptomycin (Sigma-Aldrich), and hygromycin B (Nakarai Tesque, Inc., Kyoto, Japan) were used for cell culture.

Tris (hydroxymethyl) aminomethane (FUJIFILM Wako Pure Chemicals), Opti-MEM® I Reduced Serum Medium (Thermo Fisher Scientific), plasmid DNA, Nano-Glo® Dual-Luciferase® Reporter Assay (Promega, WI, USA), HilyMax (Dojindo, Kumamoto, Japan), and hydrochloric acid (Yoneyama Yakuhin Kogyo, Osaka, Japan) were used and added with the target vector pGL4.37 [*luc2P*/ARE/Hygro] for luciferase assays.

TRIzol® (Invitrogen, MA, USA), chloroform, isopropyl alcohol (Yoneyama Yakuhin Kogyo), ethanol (FUJIFILM Wako Pure Chemicals), distilled water (Otsuka Pharmaceuticals, Tokyo, Japan), 5×PrimeScriptTM RT Master Mix, TB Green® Premix Ex Taq™ (Tli RNaseH Plus, 2 ×), and PCR primers for *HO-1* (forward 5′-TTGCCAGTGCCACCAAGTTC-3′ and reverse 5′-TCAGCAGCTCCTGCAACTCC-3′), *NQO1* (5′-GGATTGGACCGAGCTGGAA-3′ and 5′-GAAACACCCAGCCGTCAGCTA-3′), *GSTA1* (5′-TCTGCCCGTATGTCCACCTG-3′ and 5′-TGCCAACAAGGTAGTCTTGTCCA-3′), and glyceraldehyde 3-phosphate dehydrogenase (5′-GCACCGTCAAGGCTGAGAAC-3′ and 5′-TGGTGAAGACGCCAGTGGA-3′; Takara Bio Inc., Shiga, Japan) were used for RT-PCR.

### Cell culture

HepG2 cells were cultured in a RPMI 1640 medium supplemented with 10% inactivated fetal bovine serum, 100 U/mL of penicillin, and 100 µg/mL of streptomycin and stored in a CO_2_ incubator (SANYO, Tokyo, Japan) at 37°C and 5% CO_2_. The culture medium was replaced once every 3 to 4 days.

### Transfection

The cells were cultured in a 3.5 cm dish at a density of 4 × 10^5^ cells per 2 mL for 24 h. Tris (hydroxymethyl) aminomethane-hydrochloric acid buffer (20 mM, pH 7.4, 300 µL) and 4 µg of the target vector pGL4.37 [luc2P/ARE/Hygro] were added to a microtube. Then, Opti-MEM® I Reduced Serum Medium (200 µL) and 30 µL of Hilymax were added, mixed, and incubated at room temperature for 15 min. All mixture solutions were added to the culture dish medium and incubated for 4 h. Afterward, the medium was changed. After 1 week, the cells were passaged. After the cells were attached to the bottom of the dish, the medium containing 10 μg of hygromycin B per milliliter was replaced. Finally, hygromycin-resistant colonies were collected.

### Reporter assay

The transfected cells were treated with a fresh medium containing a test drug for 18 h. After removing the medium, 80 µL of fetal bovine serum and phenol red-free RPMI 1640 medium and 80 µL of One-Glo™ reagent were added to each well. After the microplate was shaken for 3 min, luminescence was measured using a microplate reader (Synergy H1, Biotek, VT, USA).

### RNA extraction

The cells were cultured in a 48-well plate at a density of 8 × 10^4^ cells per well for 24 h. The medium was replaced with a fresh medium containing the test drug and incubated for 6 h for *HO-1* RNA extraction or 18 h for *GSTA1* and *NQO1* RNA extraction. After removing the medium, 0.2 mL of TRIzol® was added to the cells. The cells were scraped to disrupt cell membranes, left for 5 min at room temperature, and collected in a microtube. Then, 40 µL of chloroform was added, and the mixture was shaken vigorously and stirred. After 5 min, each sample was centrifuged at 12,000 × g and 4°C for 15 min. Fifty microliters of the mixture was transferred from the aqueous layer to a new microtube. Then, 70 µL of isopropyl alcohol was added, and the mixture was stirred. After 10 min, each sample was centrifuged at 12,000 × g and 4°C for 10 min. After removing the supernatant, 200 µL of 75% ethanol was added, and the mixture was stirred. Each sample was centrifuged at 7,500 × g and 4°C for 5 min before the supernatant was removed. The precipitates were dissolved in 40 µL of distilled water by stirring.

### Reverse transcription

A 20 µL reverse transcription reaction consisting of 2 µL of 5 × PrimeScript RT Master Mix, 0.5 µg of total RNA, and distilled water was performed at 37°C for 15 min and 85°C for 5 s and then stored at 4°C.

### Gene expression analysis

A 25 µL real-time PCR was composed of 12.5 µL of TB Green® Premix Ex Taq™ II, 0.2 µL of 50 µM forward primer, 0.2 µL of 50 µM reverse primer, less than 100 ng of 2.0 µL template DNA, and 10.1 µL of distilled water. The PCR was performed at 95°C for 30 s, and 35 cycles of two-step PCR were performed at 95°C for 5 s and 60°C for 30 s.

The mRNA levels were quantified by calculating ΔCt and ΔΔCt values.

### Cytotoxicity assay

HepG2 cells were cultured in a 96-well plate at a density of 1 × 10^4^ cells per well for 24 h. The medium was replaced with a fresh medium containing test drugs and incubated for 48 h. Then, the medium was replaced with Cell Titer 96® diluted sixfold in a fresh medium. After 2 h, cell viability was measured at an absorbance of 490 nm.

### Statistical analysis

The results were expressed as means ± standard error. Significant differences in data were estimated using one-way analysis of variance followed by Dunnett’s multiple range test. Differences at *P* < 0.05 were considered statistically significant.

## Results

### Effects on ARE promoter activity

Oriental bezoar increased the ARE promoter activity in a dose-dependent manner, achieving a significant increase at 300 µg/mL. Similarly, bilirubin and biliverdin increased the ARE promoter activity in a dose-dependent manner, achieving a significant increase at 17.1 µM (10 µg/mL) and ≥16.2 µM (10 µg/mL), respectively. Furthermore, sulforaphane increased the ARE promoter activity at 5 µM (Fig. 1).

**Fig. 1.**
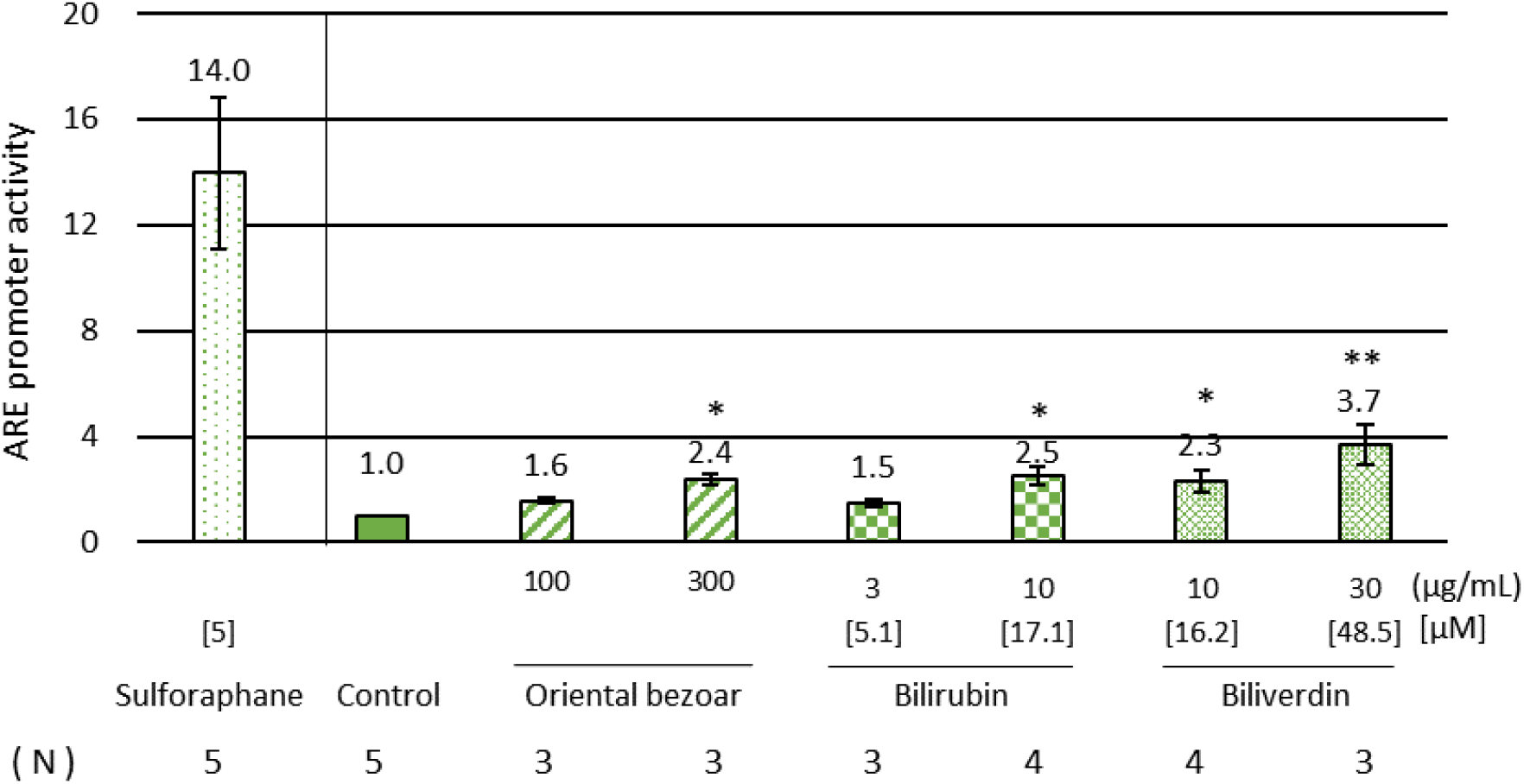
Oriental bezoar, bilirubin and biliverdin increased the ARE promoter activity in HepG2 cells. Data are expressed as mean ± SEM. *: P < 0.05, **: P < 0.01, compared to the control group using Dunnett’s test.

### Effects on the mRNA expression level of antioxidant and detoxifying enzymes

Oriental bezoar increased the levels of *HO-1* (Fig. 2), *GSTA1* (Fig. 3A), and *NQO1* (Fig. 3B) in a dose-dependent manner, resulting in significant effects at 1000 µg/mL (*HO-1*) and 1000 µg/mL (*NQO1*). Hemin increased the expression level of *HO-1* at 20 µM (Figs. 2A and 2B), whereas sulforaphane increased the expression levels of *GSTA1* (Fig. 3A) and *NQO1* (Fig. 3B) at 5 µM.

**Fig. 2.**
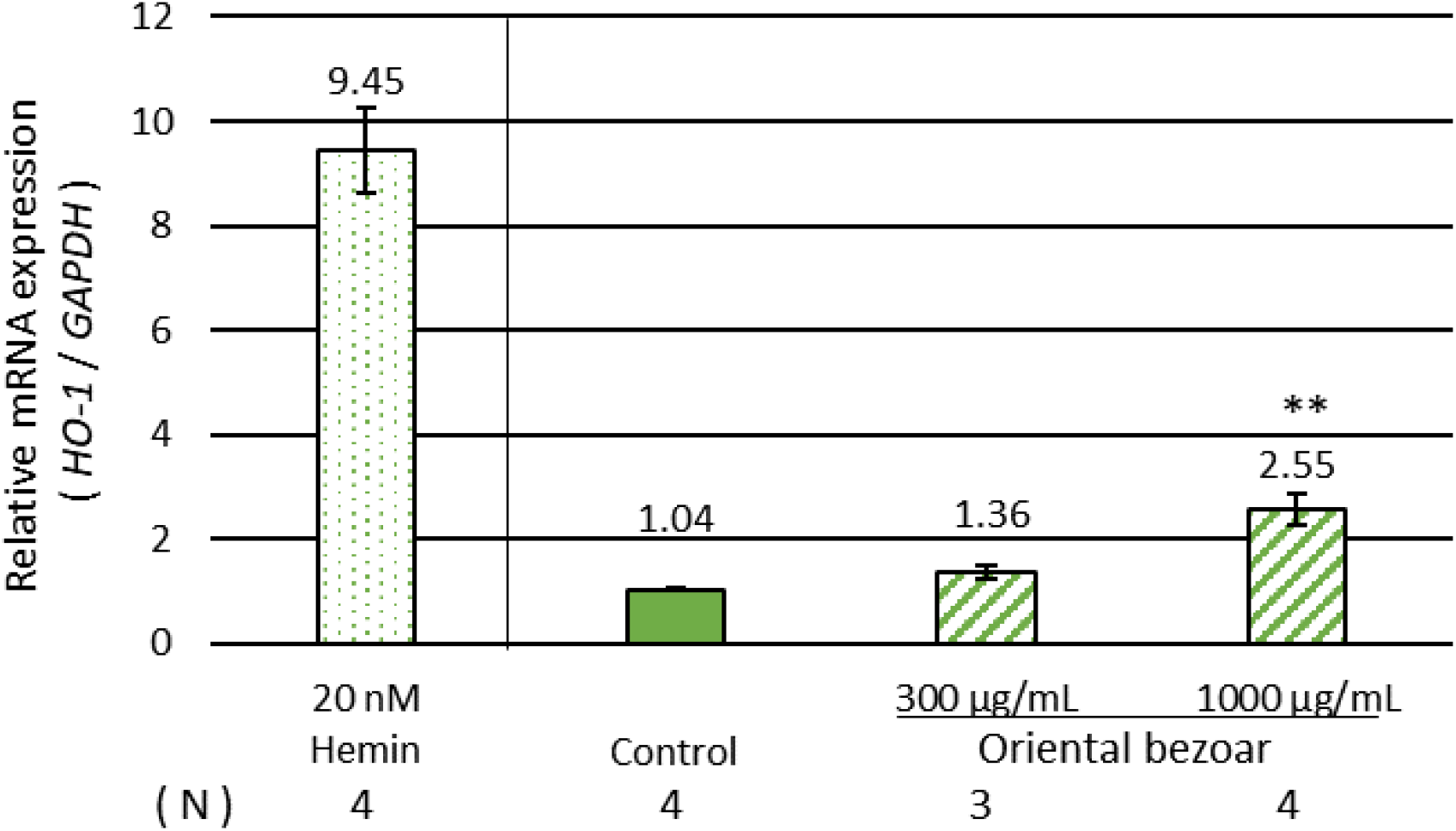
Oriental bezoar increased the expression of *HO-1* in HepG2 cells. Data are expressed as mean ± SEM. **: P < 0.01, compared to the control group using Dunnett’s test.

**Fig. 3.**
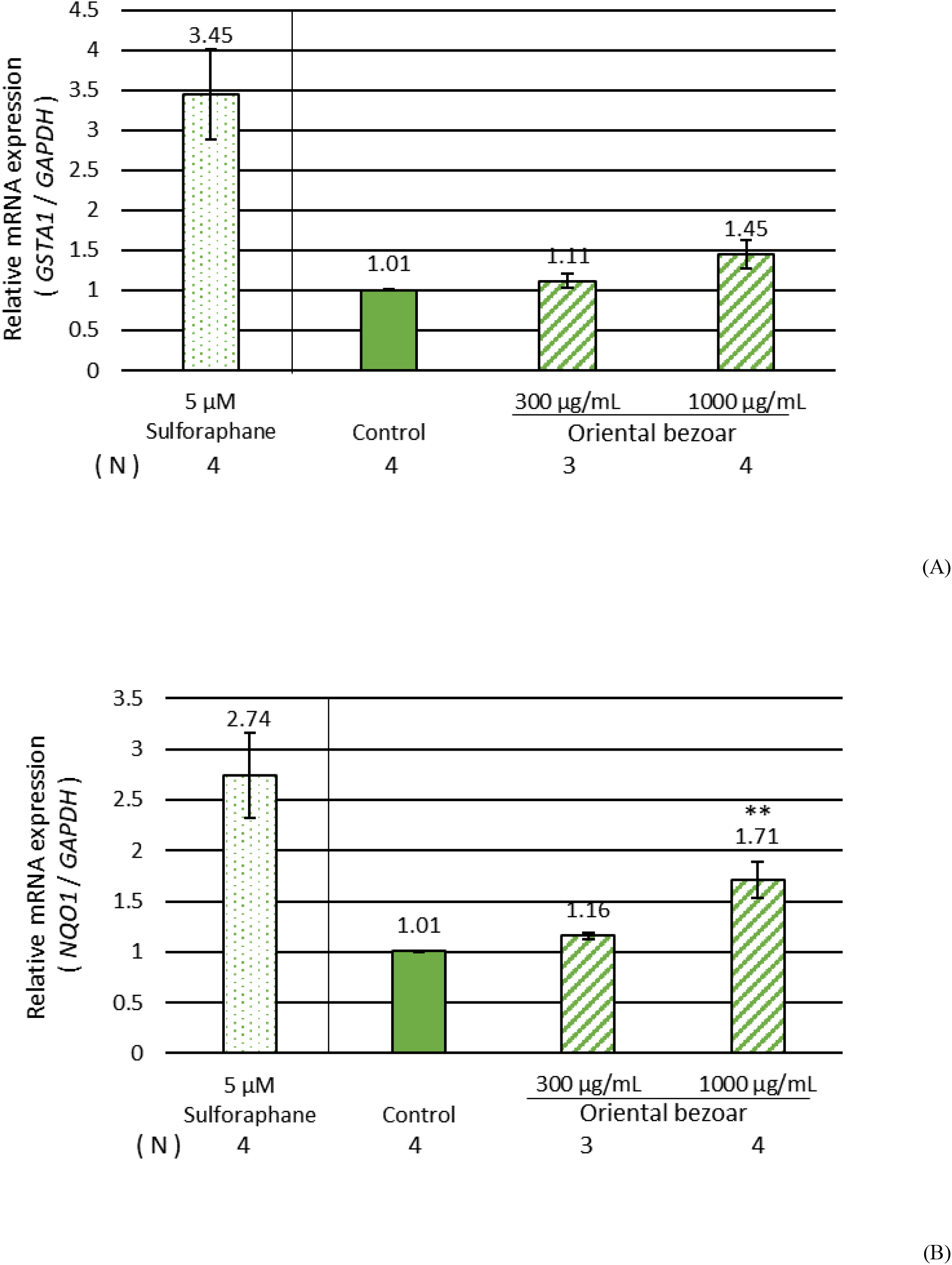
Oriental bezoar increased the expression of *GSTA1* and *NQO1* in HepG2 cells. (A) *GSTA1*, (B) *NQO1* Data are expressed as mean ± SEM. **: P < 0.01, compared to the control group using Dunnett’s test.

### Cytotoxicity

Oriental bezoar, bilirubin, and biliverdin did not exhibit cytotoxicity at 1000 µg/mL, 17.1 µM (10 µg/mL), and 48.5 µM (30 µg/mL), respectively. However, biliverdin significantly increased cell proliferation at ≥ 16.2 µM (10 µg/mL, Fig. 4).

**Fig. 4.**
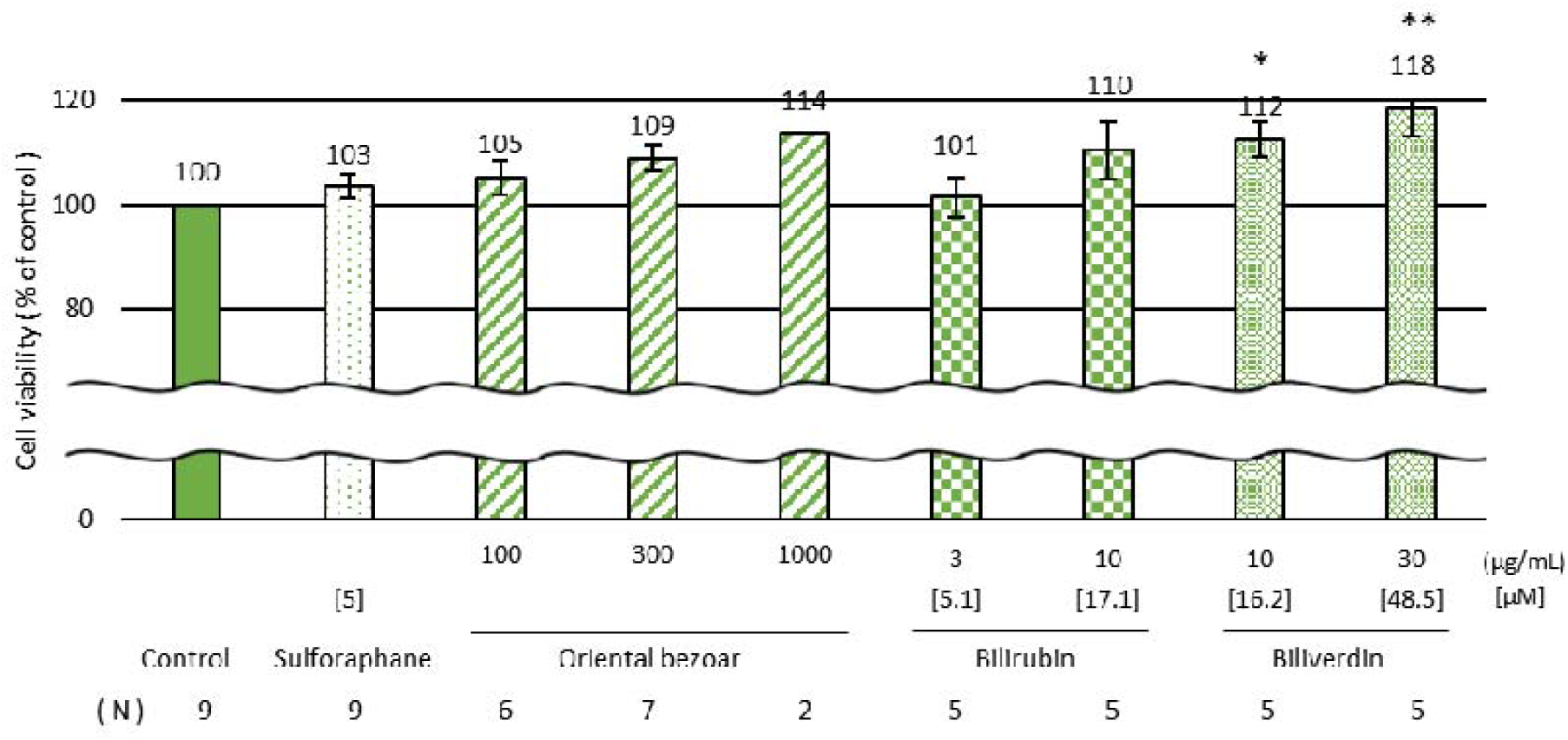
Oriental bezoar, bilirubin and biliverdin show no cytotoxicity in HepG2 cells. Data are expressed as mean ± SEM. *: P < 0.05, **: P < 0.01, compared to the control group using Dunnett’s test.

## Discussion

Oriental bezoar increased the ARE promoter activity and its downstream gene, namely, *HO-1*. HO-1 is an inducible antioxidant enzyme, which is involved in the degradation of heme into biliverdin, carbon monoxide, and free iron [10]. Biliverdin is reduced to bilirubin by biliverdin reductase [10]. Bilirubin and biliverdin have an antioxidative effect [10–12]. Based on previous reports, the activation of the Nrf2-ARE pathway could induce many antioxidant enzymes, such as γ-glutamyl cysteine ligase, thioredoxin reductase-1, superoxide dismutase, glutathione peroxidase, and catalase [13, 17]. These results indicate that oriental bezoar induces antioxidative effects via the Nrf2-ARE pathway.

Oriental bezoar could also increase the expression levels of *GSTA1* and *NQO1*, which are downstream genes of the ARE promoter. GSTA1 and NQO1 are considered as detoxifying enzymes [13, 17]. These enzymes are involved in the metabolism of carcinogens, such as benzo[a]pyrene and aflatoxin; thus, the induction of such enzymes may decrease carcinogens in the body [18]. Oriental bezoar is traditionally used for detoxification [2, 5]. Therefore, these results provide a scientific basis to support the traditional use of this crude drug.

Bilirubin and biliverdin, which are components of oriental bezoar, increased the ARE promoter activity to a certain extent. As these substances could activate the Nrf2-ARE pathway and induce *HO-1* [19], the activating effect of oriental bezoar on the Nrf2-ARE pathway is partially attributed to its constituents, namely, bilirubin and biliverdin.

However, some limitations are found in this study. The increase in expression level was only confirmed at the gene level; therefore, we must confirm this change at the protein level and the relationship between this change in gene expression levels and somatic cell or organ protection.

In summary, oriental bezoar, which is a crude drug used in oriental medicine, and its components (namely, bilirubin and biliverdin) increased the ARE promoter activity. In addition, oriental bezoar increased the mRNA expression level of *HO-1, GSTA1*, and *NQO1*. These results indicate that oriental bezoar could induce antioxidative and detoxification effects via the Nrf2-ARE pathway. However, further studies should be conducted to address the limitation.

## References

1. Shang-Hai-Ke-Xue-Ji-Shu-Chu-Ban-She. Dictionary of Chinese Materia Medica. 2nd ed. Tokyo: Shogakukan; 1985. Second volume, Bezoar; p. 787–90. Japanese.

2. State Pharmacopoeia Commission of the PRC. Pharmacopoeia of the People’s Republic of China. Beijing: ChinaMedicalScience and Technology Press; 2020. First volume, Medicinal materials, BOVIS CALCULUS; p. 72–3. Chinese.

3. Yu ZJ, Xu Y, Peng W, Liu YJ, Zhang JM, Li JS, et al. Calculus bovis: A review of the traditional usages, origin, chemistry, pharmacological activities and toxicology. J Ethnopharmacol. 2020;254:112649.

4. Wan TC, Cheng FY, Liu YT, Lin LC, Sakata R. Study on bioactive compounds of in vitro cultured Calculus Suis and natural Calculus Bovis. Anim Sci J. 2009;80(6):697–704.

5. Liu Y, Tan P, Liu S, Shi H, Feng X, Ma Q. A new method for identification of natural, artificial and in vitro cultured Calculus bovis using high-performance liquid chromatography-mass spectrometry. Pharmacogn Mag. 2015;11(42):304–10.

6. Morishita S, Saito T, Shoji M, Tanaka A, Saeki K, Ito C. [Pharmacological effects of Reiousan on experimental hepatic injuries and hepatic functions]. Folia Pharmacol Jpn. 1989;93:261–70. Japanese.

7. Morishita S, Shoji M, Oguni Y, Sugimoto C, Hirai Y, Toma S, et al. [Pharmacological studies of Reiousan which contains bezoar and ginseng: III. Effects on experimental cerebral ischemia]. Folia Pharmacol Jpn. 1991;98:435–42. Japanese.

8. Du X, Li C, Zhang S, Sun C, Zhang X, Chen C, et al. Exploring the pharmacological mechanism of calculus bovis in cerebral ischaemic stroke using a network pharmacology approach. J Ethnopharmacol. 2022;284:114507.

9. Wu T, Chang MJ, Xu YJ, Li XP, D. G, Liu D. Protective effect of Calculus Bovis Sativus on intrahepatic cholestasis in rats induced by α-naphthylisothiocyanate. Am J Chin Med. 2013;41(6):1393–405.

10. Ryter SW, Alam J, Choi AM. Heme oxygenase-1/carbon monoxide: from basic science to therapeutic applications. Physiol Rev. 2006;86(2):583–650.

11. Stocker R, Yamamoto Y, McDonagh AF, Glazer AN, Ames BN. Bilirubin is an antioxidant of possible physiological importance. Science. 1987;235(4792):1043–6.

12. Hammerman C, Goldschmidt D, Caplan MS, Kaplan M, Bromiker R, Eidelman AI, et al. Protective effect of bilirubin in ischemia-reperfusion injury in the rat intestine. J Pediatr Gastroenterol Nutr. 2002;35(3):344–9.

13. Tonelli C, Chio IIC, Tuveson DA. Transcriptional Regulation by Nrf2. Antioxid Redox Signal. 2018;29(17):1727–45.

14. Itoh K. [Disease regulation by Nrf2 antioxidant system]. SEIKAGAKU. 2009;81:447–55. Japanese.

15. Suh JH, Shenvi SV, Dixon BM, Liu H, Jaiswal AK, Liu RM, et al. Decline in transcriptional activity of Nrf2 causes age-related loss of glutathione synthesis, which is reversible with lipoic acid. Proc Natl Acad Sci U S A. 2004;101(10):3381–6.

16. Ungvari Z, Bailey-Downs L, Gautam T, Sosnowska D, Wang M, Monticone RE, et al. Age-associated vascular oxidative stress, Nrf2 dysfunction, and NF-{kappa}B activation in the nonhuman primate Macaca mulatta. J Gerontol A Biol Sci Med Sci. 2011;66(8):866–75.

17. Jung KA, Kwak MK. The Nrf2 system as a potential target for the development of indirect antioxidants. Molecules. 2010;15(10):7266–91.

18. Garg R, Gupta S, Maru GB. Dietary curcumin modulates transcriptional regulators of phase I and phase II enzymes in benzo[a]pyrene-treated mice: mechanism of its anti-initiating action. Carcinogenesis. 2008;29(5):1022–32.

19. Huang Y, Li J, Li W, Ai N, Jin H. Biliverdin/Bilirubin Redox Pair Protects Lens Epithelial Cells against Oxidative Stress in Age-Related Cataract by Regulating NF-κB/iNOS and Nrf2/HO-1 Pathways. Oxid Med Cell Longev. 2022;2022:7299182.

